# The Diagnostic Potential & Interactive Dynamics of the Colorectal Cancer Virome

**DOI:** 10.1101/152868

**Authors:** Geoffrey D Hannigan, Melissa B Duhaime, Mack T Ruffin, Charlie C Koumpouras, Patrick D Schloss

**Affiliations:** Department of Microbiology & Immunology, University of Michigan, Ann Arbor, Michigan, 48109; Department of Ecology and Evolutionary Biology, University of Michigan, Ann Arbor, Michigan, 48109; Department of Family and Community Medicine, Pennsylvania State University Hershey Medical Center, Hershey, Pennsylvania, 17033

## Abstract

Human viruses (those that infect human cells) have been associated with many cancers, largely due to their mutagenic and functionally manipulative abilities. Despite this, cancer microbiome studies have almost exclusively focused on bacteria instead of viruses. We began evaluating the cancer virome by focusing on colorectal cancer, a primary cause of morbidity and mortality throughout the world, and a cancer linked to altered colonic bacterial community compositions but with an unknown association with the gut virome. We used 16S rRNA gene, whole shotgun metagenomic, and purified virus metagenomic sequencing of stool to evaluate the differences in human colorectal cancer virus and bacterial community composition. Through random forest modeling we identified differences in the healthy and colorectal cancer virome. The cancer-associated virome consisted primarily of temperate bacteriophages that were also predicted to be bacteria-virus community network hubs. These results provide foundational evidence that bacteriophage communities are associated with colorectal cancer and potentially impact cancer progression by altering the bacterial host communities.

## Importance

Colorectal cancer is a leading cause of cancer-related death in the United States and worldwide. Its risk and severity have been linked to colonic bacterial community composition. Although human-specific viruses have been linked to other cancers and diseases, little is known about colorectal cancer virus communities. We addressed this knowledge gap by identifying differences in colonic virus communities in the stool of colorectal cancer patients and how they compared to bacterial community differences. The results suggested an indirect role for the virome in impacting colorectal cancer by modulating their associated bacterial community. These findings both support a biological role for viruses in colorectal cancer and provide a new understanding of basic colorectal cancer etiology.

## Introduction

The human gut virome is the community of all viruses found in the gut, including bacteriophages (viruses that only infect bacteria), eukaryotic viruses (viruses that only infect eukaryotic cells), and human-specific viruses (viruses that only infect human cells). Due to their mutagenic abilities and propensity for functional manipulation, human viruses are strongly associated with, and in many cases cause, cancer (1–4). Because bacteriophages are crucial for bacterial community stability and composition (5–7), and members of those bacterial communities have been implicated as oncogenic agents (8–11), bacteriophages have the potential to indirectly impact cancer as well. The gut virome therefore has a potential to be associated with, and potentially impact, human cancer. Altered human virome composition and diversity have already been identified in diseases including periodontal disease (12), HIV (13), cystic fibrosis (14), antibiotic exposure (15, 16), urinary tract infections (17), and inflammatory bowel disease (18). The strong association of bacterial communities with colorectal cancer, the previous identification of human-specific viruses that cause cancer, and the precedent for the virome to impact other human diseases suggest that colorectal cancer may be associated with altered virus communities.

Colorectal cancer is the second leading cause of cancer-related deaths in the United States (19). The US National Cancer Institute estimates over 1.5 million Americans were diagnosed with colorectal cancer in 2016 and over 500,000 Americans died from the disease (19). Growing evidence suggests that an important component of colorectal cancer etiology may be perturbations in the colonic bacterial community (8, 10, 11, 20, 21). Work in this area has led to a proposed disease model in which bacteria colonize the colon, develop biofilms, promote inflammation, and enter an oncogenic synergy with the cancerous human cells (22). This association also has allowed researchers to leverage bacterial community signatures as biomarkers to provide accurate, noninvasive colorectal cancer detection from stool (8, 23, 24). While an understanding of colorectal cancer bacterial communities has proven fruitful both for disease classification and for identifying the underlying disease etiology, bacteria are only a subset of the colon microbiome. Viruses are another important component of the colon microbial community that have yet to be studied in the context of colorectal cancer. We evaluated disruptions in virus and bacterial community composition in a human cohort whose stool was sampled at the three relevant stages of cancer development: healthy, adenomatous, and cancerous.

Colorectal cancer progresses in a stepwise process that begins when healthy tissue develops into a precancerous polyp (i.e., adenoma) in the large intestine (25). If not removed, the adenoma may develop into a cancerous lesion that can invade and metastasize, leading to severe illness and death. Progression to cancer can be prevented when precancerous adenomas are detected and removed during routine screening (26, 27). Survival for colorectal cancer patients may exceed 90% when the lesions are detected early and removed (26). Thus, work that aims to facilitate early detection and prevention of progression beyond early cancer stages has great potential to inform therapeutic development.

Here we begin to address the knowledge gap of whether virus community composition is altered in colorectal cancer and, if it is, how those differences might impact cancer progression and severity. We also aimed to evaluate the virome’s potential for use as a diagnostic biomarker. The implications of this study are threefold. *First*, this work supports a biological role for the virome in colorectal cancer development and suggests that more than the bacterial members of the associated microbial communities are involved in the process. *Second*, we present a supplementary virus-based approach for classification modeling of colorectal cancer using stool samples. *Third*, we provide initial support for the importance of studying the virome as a component of the microbiome ecological network, especially in cancer.

## Results

### Sample Collection and Processing

Our study cohort consisted of stool samples collected from 90 human subjects, 30 of whom had healthy colons, 30 of whom had adenomas, and 30 of whom had carcinomas **(Figure 1)**. Half of each stool sample was used to sequence the bacterial communities using both 16S rRNA gene and shotgun sequencing techniques. The 16S rRNA gene sequencing was performed for a previous study, and the sequences were re-analyzed using contemporary methods (8). The other half of each stool sample was purified for virus like particles (VLPs) before genomic DNA extraction and shotgun metagenomic sequencing. In the VLP purification, cells were disrupted and extracellular DNA degraded **(Figure 1)** to allow the exclusive analysis of viral DNA within virus capsids. In this manner, the *extracellular virome* of encapsulated viruses was targeted.

**Figure 1:**
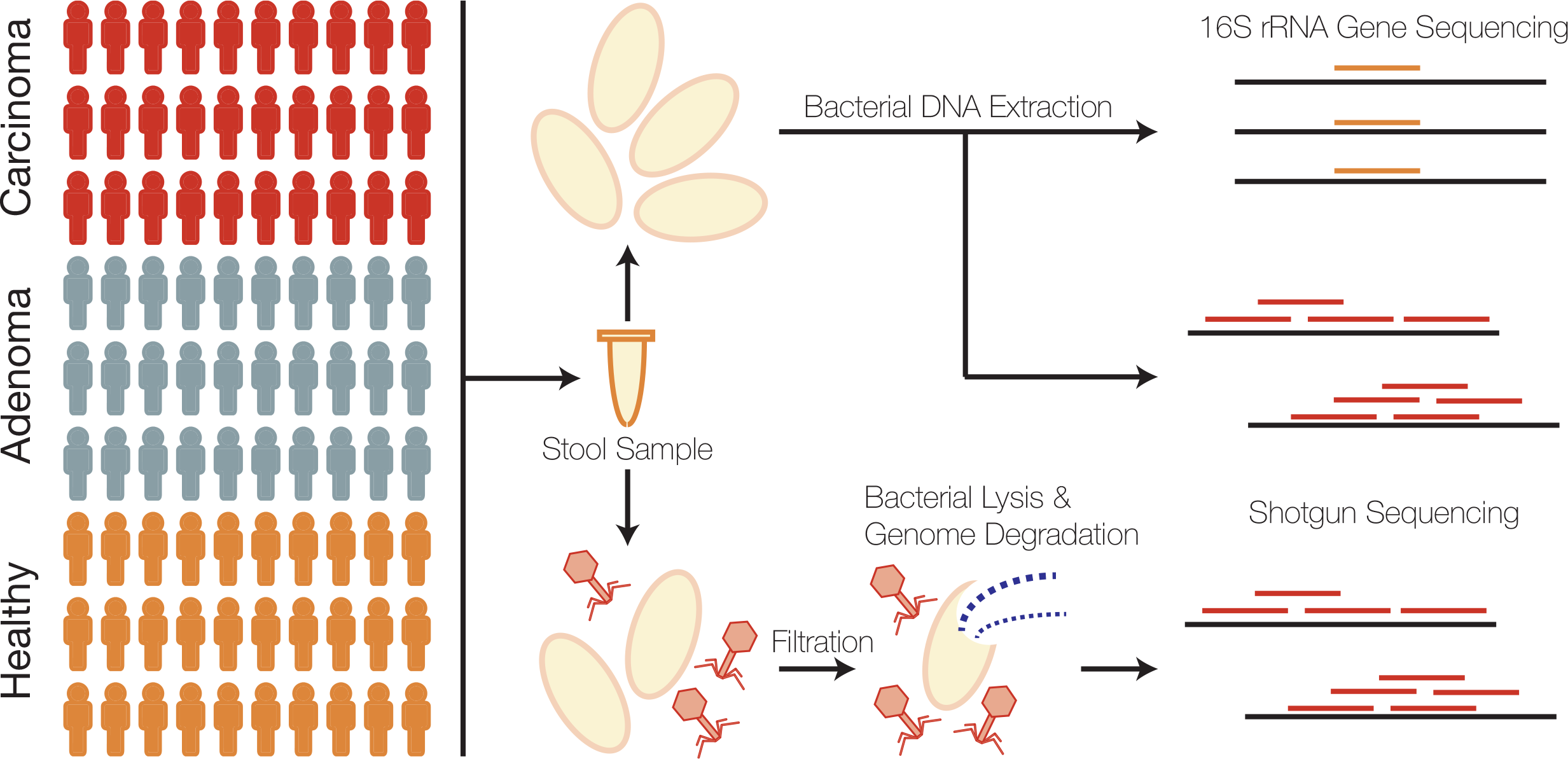
Cohort and sample processing outline. Thirty subject stool samples were collected from healthy, adenoma (pre-cancer), and carcinoma (cancer) patients. Stool samples were split into two aliquots, the first of which was used for bacterial sequencing and the second which was used for virus sequencing. Bacterial sequencing was done using both 16S rRNA amplicon and whole metagenomic shotgun sequencing techniques. Virus samples were purified for viruses using filtration and a combination of chloroform (bacterial lysis) and DNase (exposed genomic DNA degradation). The resulting encapsulated virus DNA was sequenced using whole metagenomic shotgun sequencing.

Each extraction was performed with a blank buffer control to detect contaminants from reagents or other unintentional sources. Only one of the nine controls contained detectable DNA at a minimal concentration of 0. 011 ng/*μ*l, thus providing evidence of the enrichment and purification of VLP genomic DNA over potential contaminants **(Figure S1 A)**. As expected, these controls yielded few sequences and were almost entirely removed while rarefying the datasets to a common number of sequences **(Figure S1 B)**. The high quality phage and bacterial sequences were assembled into highly covered contigs longer than 1 kb **(Figure S2)**. Because contigs represent genome fragments, we further clustered related bacterial contigs into operational genomic units (OGUs) and viral contigs into operational viral units (OVUs) (**Figure S2** **- S3)** to approximate organismal units.

### Unaltered Diversity in Colorectal Cancer

Microbiome and disease associations are often described as being of an altered diversity (i.e. “dysbiotic”). Therefore, we first evaluated the influence of colorectal cancer on virome OVU diversity. We evaluated differences in communities between disease states using the Shannon diversity, richness, and Bray-Curtis metrics. We observed no significant alterations in either Shannon diversity or richness in the diseased states as compared to the healthy state **(Figure S4 C-D)**. There was no statistically significant clustering of the disease groups (ANOSIM p-value = 0.6, **Figure S4**). Notably, there was a significant difference between the few blank controls that remained after rarefying the data and the other study groups (ANOSIM p-value < 0.001, **Figure S5)**, further supporting the quality of the sample set. In summary, standard alpha and beta diversity metrics were insufficient for capturing virus community differences between disease states **(Figure S4)**. This is consistent with what has been observed when the same metrics were applied to 16S rRNA gene sequences and metagenomic samples (8, 23, 24) and points to the need for alternate approaches to detect the impact of colorectal cancer disease state on these community structures.

### Virome Composition in Colorectal Cancer

As opposed to the diversity metrics discussed above, OTU-based relative abundance profiles generated from 16S rRNA gene sequences are effective for classifying stool samples as originating from individuals with healthy, adenomatous, or cancerous colons (8, 23). By using classification models instead of attempting to identify single differentially abundant OTUs, these and other studies have been successful in capturing complex community relationships in which differences in taxonomic relative abundance are considered in the context of other taxa. The exceptional performance of bacteria in these classification models supports a role for bacterial functionality in colorectal cancer. We built off of these findings by evaluating the ability of virus community signatures to classify stool samples and compared their performance to models built using bacterial community signatures.

To identify the altered virus communities associated with colorectal cancer, we built and tested random forest models for classifying stool samples as belonging to individuals with either cancerous or healthy colons. We confirmed that our bacterial 16S rRNA gene model replicated the performance of the original report which used logit models instead of random forest models **(Figure 2 A)** (8). We then compared the bacterial OTU model to a model built using OVU relative abundances. The viral model performed as well as the bacterial model (corrected p-value = 0.6), with the viral and bacterial models achieving mean area under the curve (AUC) values of 0.768 and 0.775, respectively **(Figure 2 A - B)**. To evaluate the ability of both bacterial and viral biomarkers to classify samples, we built a combined model that used both bacterial and viral community data. The combined model did not yield a statistically significant performance improvement beyond the viral (corrected p-value = 0.08) and bacterial (corrected p-value = 0.1) models, yielding an AUC of 0.807 **(Figure 2 A - B)**.

**Figure 2:**
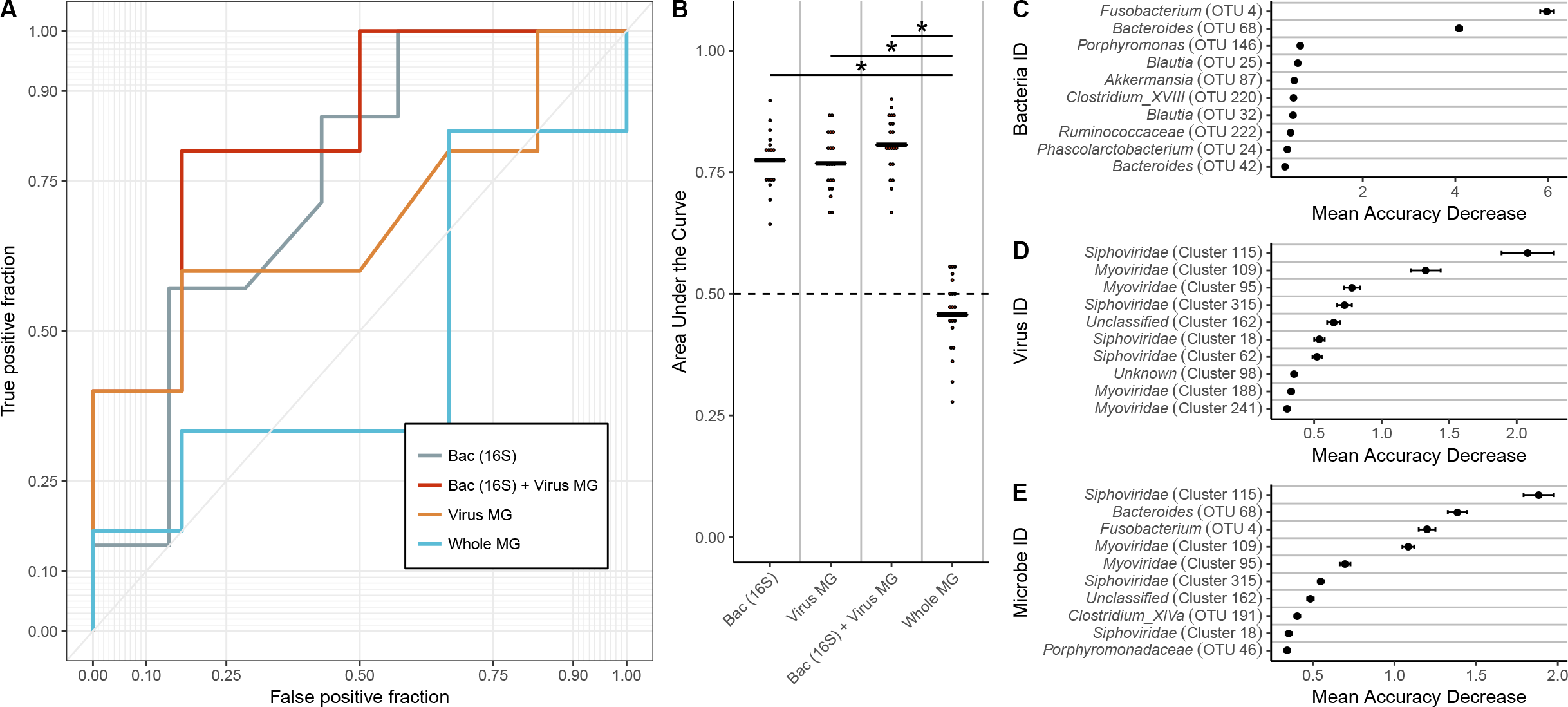
Results from healthy vs cancer classification models built using virome signatures, bacterial 16S rRNA gene sequence signatures, whole metagenomic signatures, and a combination of virome and 16S rRNA gene sequence signatures. A) An example ROC curve for visualizing the performance of each of the models for classifying stool as coming from either an individual with a cancerous or healthy colon. B) Quantification of the AUC variation for each model, and how it compared to each of the other models based on 15 iterations. A pairwise Wilcoxon test with a false discovery rate multiple hypothesis correction demonstrated that all models are significantly different from each other (p-value < 0.01). C) Mean decrease in accuracy (measurement of importance) of each operational taxonomic unit within the 16S rRNA gene classification model when removed from the classification model. Mean is represented by a point, and bars represent standard error. D) Mean decrease in accuracy of each operational virus unit in the virome classification model. E) Mean decrease in accuracy of each operational genomic unit and operational taxonomic unit in the model using both 16S rRNA gene and virome features.

We compared viral metagenomic methods to bacterial metagenomic methods by building a viral model and a model built using OGU relative abundance profiles from bacterial metagenomic shotgun sequencing data. This bacterial model performed worse than the other models (mean AUC = 0.458) **(Figure 2 A - B)**. To determine the cause of the discrepancy between the two bacterial sequencing methods, we attempted to compare the approaches at a common sequencing depth. This revealed that the bacterial 16S rRNA gene model was strongly driven by sparse and low abundance OTUs **(Figure S6)**. Removal of OTUs with a median abundance of zero resulted in the removal of six OTUs, and a loss of model performance down to what was observed in the metagenome-based model **(Figure S6 A)**. The majority of these OTUs had a relative abundance lower than 1% across the samples **(Figure S6 B)**. Although the features in the viral model also were of low abundance **(Figure S8F)**, the coverage was sufficient for high model performance, likely because viral genomes are orders of magnitude smaller than bacterial genomes.

The association between the bacterial and viral communities and colorectal cancer was driven by a few important microbes. *Fusobacterium* was the primary driver of the bacterial association with colorectal cancer, which is consistent with its previously described oncogenic potential **(Figure 2 C)**(22). The virome signature also was driven by a few OVUs, suggesting a role for these viruses in tumorigenesis **(Figure 2 D)**. It is also important to note that while these viruses were driving the signature, the magnitude of their importance and the significance of those values was noticibly less than the bacterial 16S signature, suggesting that unlike what is observed in the bacteria, there are many viruses that are associated with the cancerous state. The identified viruses were bacteriophages, belonging to *Siphoviridae*, *Myoviridae*, and phage taxa that could not be confidently identified beyond their broad phage identification (i.e. “unclassified”). Viruses, which were confirmed to not have genomic similarity to known bacterial genomes, were unidentifiable (denoted “unknown”). This is common in viromes across habitats; studies have reported as much as 95% of virus sequences belonging to unknown genomic units (14, 28–30). When the bacterial and viral community signatures were combined, both bacterial and viral organisms drove the community association with cancer **(Figure 2 E)**.

### Phage Influence Between CRC Stages

Because previous work has identified shifts in which bacteria were most important at different stages of colorectal cancer (8, 20, 22), we explored whether shifts in the relative influence of phages could be detected between healthy, adenomatous, and cancerous colons. We evaluated community shifts between the disease stage transitions (healthy to adenomatous and adenomatous to cancerous) by building random forest models to compare only the diagnosis groups around the transitions. While bacterial OTU models performed equally well for all disease class comparisons, the virome model performances differed **(Figure S7 A-B)**. Like bacteria **(Figure S7 F-H)**, different virome members were important between the healthy to adenomatous and adenomatous to cancerous stages **(Figure S7C-E)**.

After evaluating our ability to classify samples between two disease states, we performed a three-class random forest model including all disease states. The 16S rRNA gene model yielded a mean AUC of 0.784 and outperformed the viral community model, which yielded a mean AUC of 0.654 (p-value < 0.001, **Figure S8 A-C**). The microbes important for the healthy versus cancer and healthy versus adenoma models were also important for the three-class model **(Figure S8 D-E)**. The most important bacterium in the two and three class models was the same *Fusobacterium* (OTU 4) (**Figure 2 C**, **Figure S8 D)**. The viruses most important to the three-class model were identified as bacteriophages (**Figure 2 D**, **Figure S8 E)**, but not all important OVUs were of increased abundance in the diseased state **(Figure S8 F)**.

### Phage Dominance in CRC Virome

Differences in the colorectal cancer virome could have been driven by eukaryotic (human) viruses or by bacteriophages. To better understand the types of viruses that were important for colorectal cancer, we identified the virome OVUs as being similar to either eukaryotic viruses or bacteriophages. The most important viruses to the classification model were identified as bacteriophages (**Figure S8)**. Overall, we were able to identify 78.8% of the OVUs as known viruses, and 93.8% of those viral OVUs aligned to bacteriophage reference genomes. It is important to note that this could have been influenced by our methodological biases against enveloped viruses (more common of eukaryotic viruses than bacteriophage), due to chloroform and DNase treatment for purification.

We evaluated whether the phages in the community were primarily lytic (i.e. obligately lyse their hosts after replication) or temperate (i.e. able to integrate into their host’s genome to form a lysogen, and subsequently transition to a lytic mode). We accomplished this by identifying three markers for temperate phages in the OVU representative sequences: 1) presence of phage integrase genes, 2) presence of known prophage genes, according the the ACLAME (ACLAssification of Mobile genetic Elements) database, and 3) nucleotide similarity to regions of bacterial genomes (29, 31, 32). We found that the majority of the phages were temperate and that the overall fraction of temperate phages remained consistent throughout the healthy, adenomatous, and cancerous stages **(Figure 3)**. These findings were consistent with previous reports suggesting the gut virome is primarily composed of temperate phages (13, 18, 31, 33).

**Figure 3:**
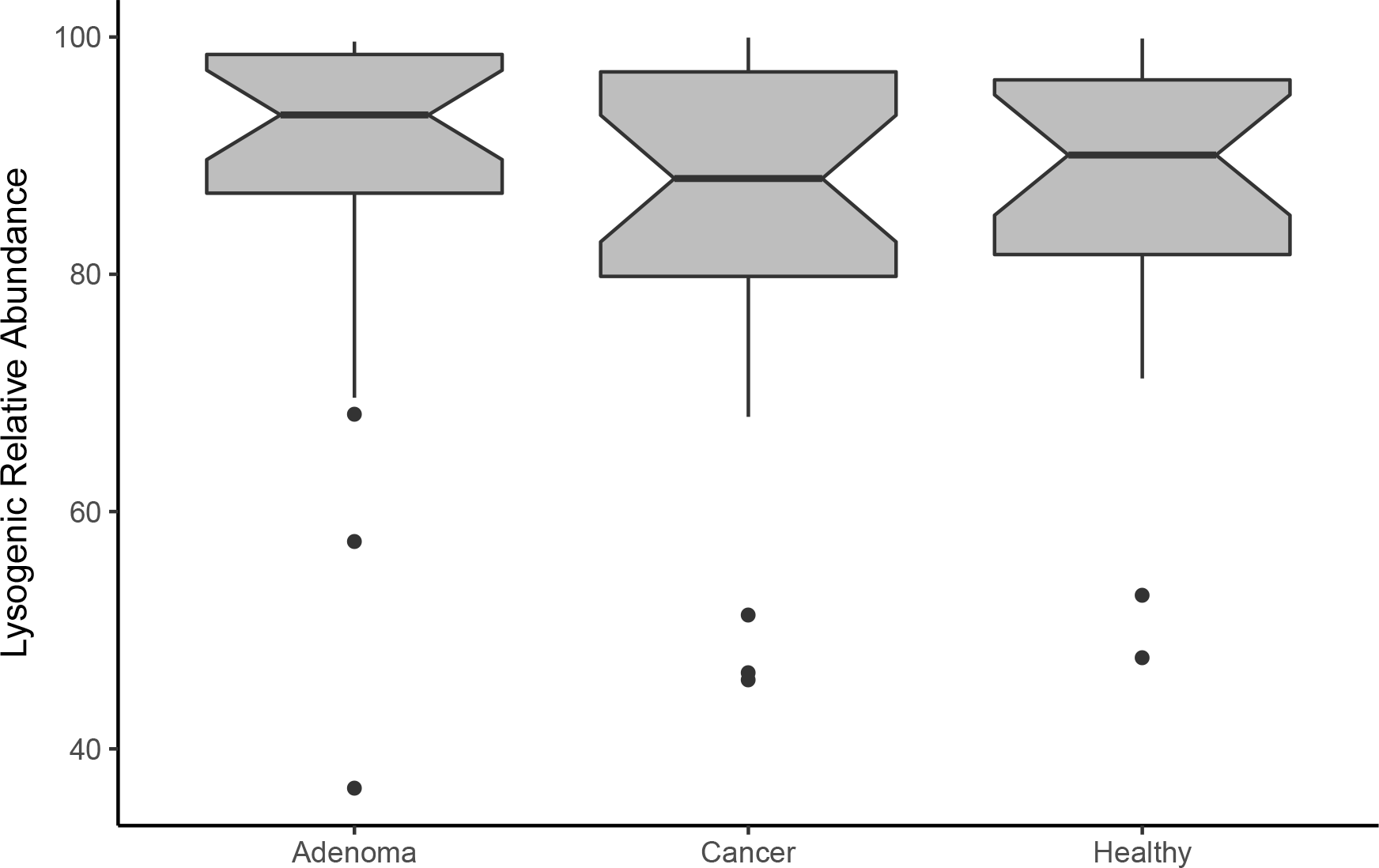
Lysogenic phage relative abundance in disease states. Phage OVUs were predicted to be either lytic or lysogenic, and the relative abundance of lysogenic phages was quantified and represented as a boxplot. No disease groups were statistically significant.

### Community Context of Influential Phages

Because the link between colorectal cancer and the virome was driven by bacteriophages (as opposed to non-bacterial viruses), we tested a potential hypothesis that the virome signal was a mere reflection of the bacterial signal, and thus highly correlated with the bacterial signal. If this hypothesis were true, we would expect a correlation between the relative abundances of influential bacterial OTUs and virome OVUs. Instead, we observed a strikingly low correlation between bacterial and viral relative abundances **(Figure 4 A,C)**. Overall, there was an absence of correlation between the most influential OVUs and bacterial OTUs **(Figure 4 B)**. This evidence supported our null hypothesis that the influential viral OVUs were not primarily reflections of influential bacteria.

**Figure 4:**
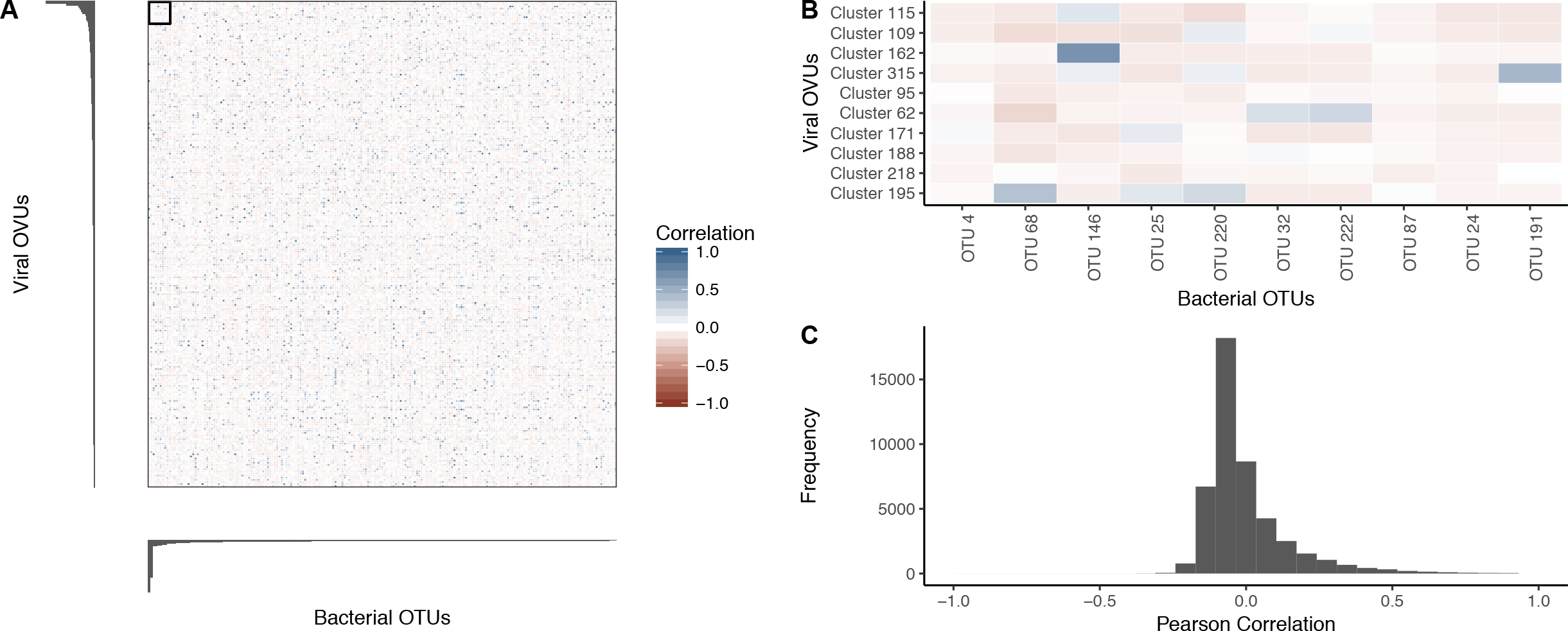
Relative abundance correlations between bacterial OTUs and virome OVUs. A) Pearson correlation coefficient values between all bacterial OTUs (x-axis) and viral OVUs (y-axis) with blue being positively correlated and red being negatively correlated. Bar plots indicate the viral (left) and bacterial (bottom) operational unit importance in their colorectal cancer classification models, such that the most important units are in the top left corner. B) Magnification of the boxed region in panel (A), highlighting the correlation between the most important bacterial OTUs and virome OVUs. The most important operational units are in the top left corner of the heatmap, and the correlation scale is the same as panel (A). C) Histogram quantifying the frequencies of Pearson correlation coefficients between all bacterial OTUs and virome OVUs.

Given these findings, we posited that the most influential phages were acting by infecting a wide range of bacteria in the overall community, instead of just the influential bacteria. In other words, we hypothesized that the influential bacteriophages were community hubs (i.e. central members) within the bacteria and phage interactive network. We investigated the potential host ranges of all phage OVUs using a previously developed random forest model that relies on sequence features to predict which phages infected which bacteria in the community **(Figure 5A)** (34). The predicted interactions were then used to identify phage community hubs. We calculated the alpha centrality (i.e. measure of importance in the ecological network) of each phage OVU’s connection to the rest of the network. The phages with high centrality values were defined as community hubs. Next, the centrality of each OVU was compared to its importance in the colorectal cancer classification model. Phage OVU centrality was significantly and positively correlated with importance to the disease model (p-value = 0.004, R = 0.176), suggesting that phages that were important in driving colorectal cancer also were more likely to be community hubs **(Figure 5 B)**. Together these findings supported our hypothesis that influential phages were hubs within their microbial communities and had broad host ranges.

**Figure 5:**
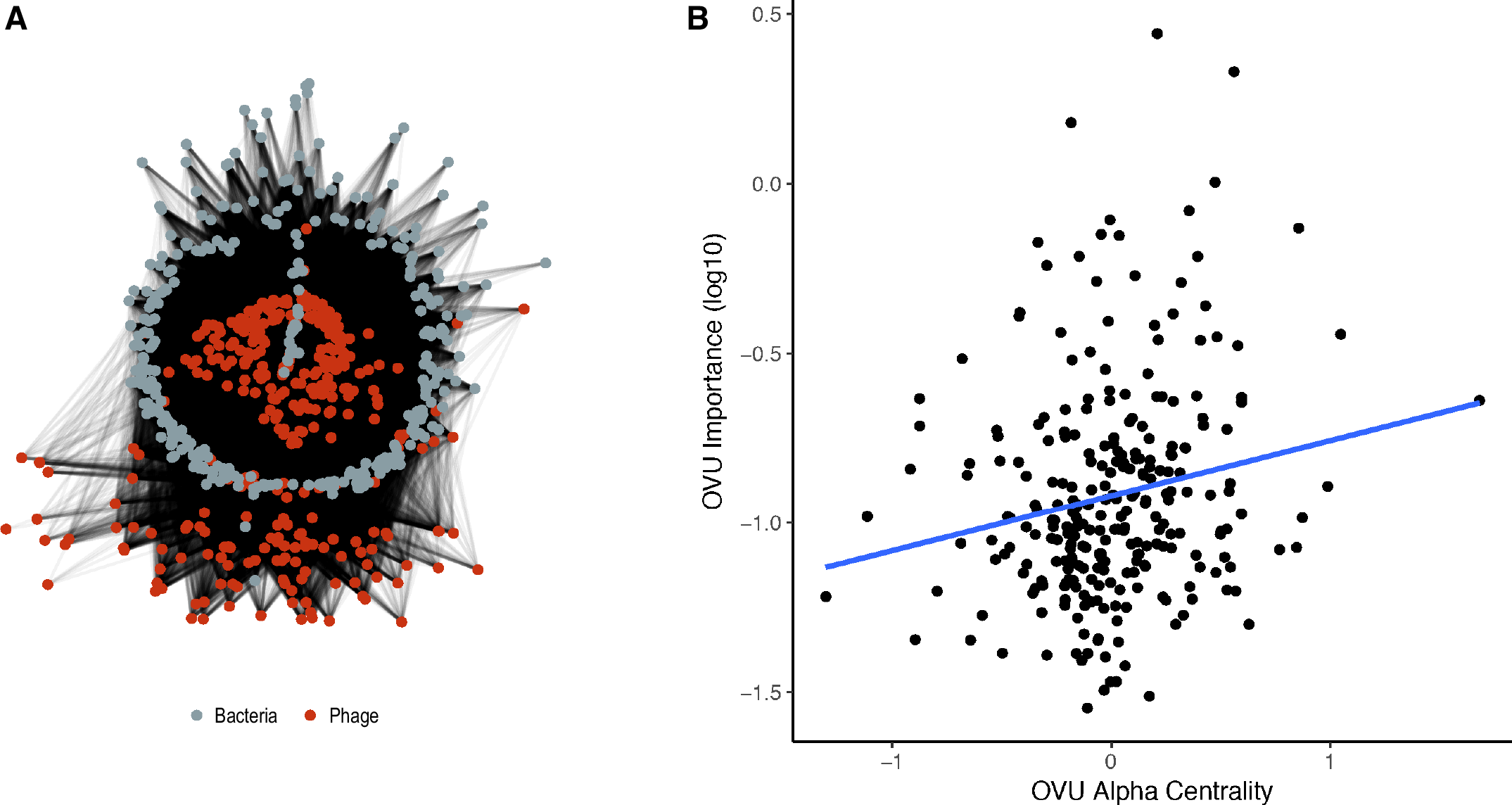
Community network analysis utilizing predicted interactions between bacteria and phage operational genomic units. A) Visualization of the community network for our colorectal cancer cohort. B) Scatter plot illustrating the correlation between importance (mean decrease in accuracy) and the degree of centrality for each OVU. A linear regression line was fit to illustrate the correlation (blue) which was found to be statistically significantly and weakly correlated (p-value = 0.00409, R = 0.176).

## Discussion

Because of their propensity for mutagenesis and capacity for modulating their host functionality, many human viruses are oncogenic (1–4). Some bacteria also have oncogenic properties, suggesting that bacteriophages, a component of the human virome in addition to human-specific viruses, may play an indirect role in promoting carcinogenesis by influencing bacterial community composition and dynamics (8–10). Despite their carcinogenic potential and the strong association between bacteria and colorectal cancer, a link between virus colorectal communities and colorectal cancer has yet to be evaluated. Here we show that, like colonic bacterial communities, the colon virome was altered in patients with colorectal cancer relative to those with healthy colons. Our findings support a working hypothesis for oncogenesis by phage-modulated bacterial community composition.

Based on our findings, we have developed a conceptual model to be tested in our future studies aimed at elucidating the role the colonic virome plays in colorectal cancer **(Figure 6 A)**. We found that basic diversity metrics of alpha diversity (richness and Shannon diversity) and beta diversity (Bray-Curtis dissimilarity) were insufficient for identifying virome community differences between healthy and cancerous states. By implementing a machine learning approach (random forest classification) to leverage inherent, complex patterns not detected by diversity measures, we were able to detect strong associations between the colon virus community composition and colorectal cancer. The dsDNA virome of colorectal cancer was composed primarily of bacteriophages. These phage communities were not exclusively predators of the most influential bacteria, as demonstrated by the lack of correlation between the abundances of the bacterial and phage populations. Instead, we identified influential phages as being community hubs, suggesting phages influence cancer by altering the greater bacterial community instead of directly modulating the influential bacteria. Our previous work has shown that modifying colon bacterial communities alters colorectal cancer progression and tumor burden in mice (10, 20). This provides a precedent for phages indirectly influencing colorectal cancer progression by altering the bacterial community composition. Overall, our data support a model in which the bacteriophage community modulates the bacterial community, and through those interactions indirectly influences the bacteria driving colorectal cancer progression **(Figure 6A)**. Although our evidence suggested phages indirectly influenced colorectal cancer development, we were not able to rule out the role of phages directly interacting with the human host (35, 36).

**Figure 6:**
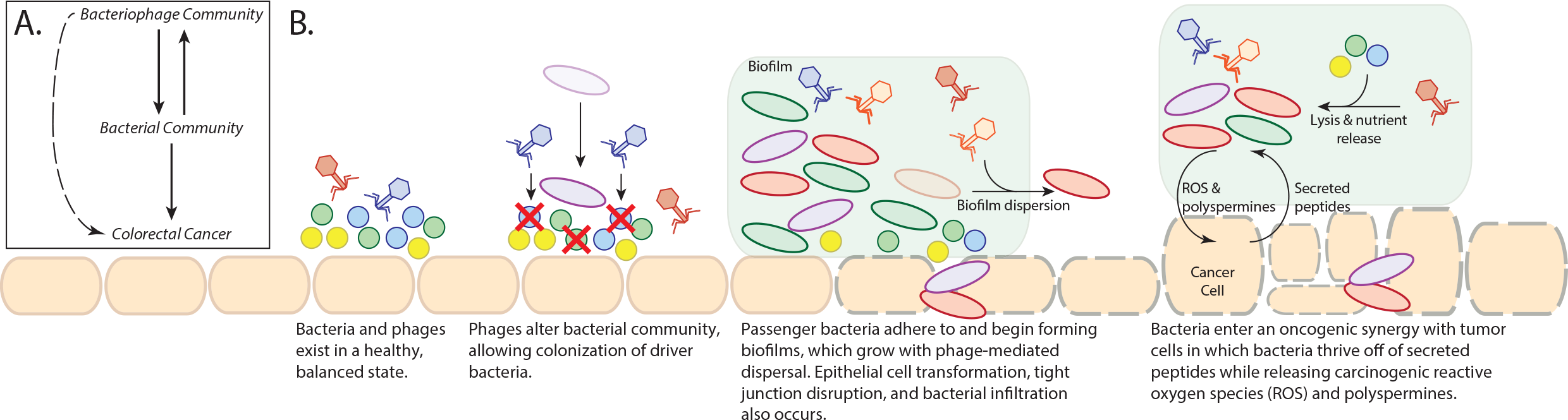
Final working hypothesis from this study. These panels summarize our thoughts on our results and represent interesting future directions that we predict will build on the presented work. A) Basic model illustrating the connections between the virome, bacterial communities, and colorectal cancer. B) Working hypothesis of how the bacteriophage community is associated with colorectal cancer and the associated bacterial community.

In addition to modeling the potential connections between virus communities, bacterial communities, and colorectal cancer, we also used our data and existing knowledge of phage biology to develop a working hypothesis for the mechanisms by which this may occur. This was done by incorporating our findings into a current model for colorectal cancer development **(Figure 6 B)** (22), although it is important to note that there are also many other alternative hypotheses by which the system could be operating. We hypothesize that the process begins with broadly infectious phages in the colon lysing, and thereby disrupting, the existing bacterial communities. This shift opens novel niche space that enabled opportunistic bacteria (such as *Fusobacterium nucleatum)* to colonize. Once the initial influential founder bacteria establish themselves in the epithelium, secondary opportunistic bacteria are able to adhere to the founders, colonize, and establish a biofilm. Phages may play a role in biofilm dispersal and growth by lysing bacteria within the biofilm, a process important for effective biofilm growth (37). The oncogenic bacteria may then be able to transform the epithelial cells and disrupt tight junctions to infiltrate the epithelium, thereby initiating an inflammatory immune response. As the adenomatous polyps developed and progressed towards carcinogenesis, we observed a shift in the phages and bacteria whose relative abundances were most influential. As the bacteria enter their oncogenic synergy with the epithelium, we conjecture that the phages continue mediating biofilm dispersal. This process would thereby support the colonized oncogenic bacteria by lysing competing cells and releasing nutrients to other bacteria in the form of cellular lysates. In addition to highlighting the likely mechanisms by which the colorectal cancer virome is interacting with the bacterial communities this model will guide future research investigations of the role the virome plays colorectal cancer.

Our working hypothesis represents a conceptualization of areas for future work, which will be required to characterize the colorectal cancer microbiome at the functional, mechanistic level. There are many different ways in which this system may operating, and our working hypothesis is one. For example, it is possible that the bacterial communities cause a change in the virome instead of the virome altering the bacterial communities. To better understand this system, future studies will include larger cohort human studies, further *in vitro* and *in vivo* mechanistic experimentation, and attempts at community studies using absolute abundance values instead of relative abundance, which would allow for more accurate community dynamic modeling. Overall, this study provides a conceptual foundation to direct future characterization of the colorectal cancer microbiome at the functional, mechanistic level.

In addition to the diagnostic ramifications for understanding the colorectal cancer microbiome, our findings suggest that viruses, while understudied and currently under-appreciated in the human microbiome, are likely to be an important contributor to human disease. Viral community dynamics have the potential to provide an abundance of information to supplement those of bacterial communities. Evidence has suggested that the virome is a crucial component to the microbiome and that bacteriophages are important players. Bacteriophage and bacterial communities cannot maintain stability and co-evolution without one another (6, 38). Not only is the human virome an important element to consider in human health and disease (12–18), but our findings support that it is likely to have a significant impact on cancer etiology and progression.

## Materials and Methods

### Analysis Source Code & Data Availability

All study sequences are available on the NCBI Sequence Read Archive under the BioProject ID `PRJNA389927`.

All associated source code is available at the following GitHub repository: https://github.com/SchlossLab/Hannigan_CRCVirome_mBio_2018

### Study Design and Patient Sampling

This study was approved by the University of Michigan Institutional Review Board and all subjects provided informed consent. Design and sampling of this sample set have been reported previously (8). Briefly, whole evacuated stool was collected from patients who were 18 years of age or older, able to provide informed consent, have had colonoscopy and histologically confirmed colonic disease status, had not had surgery, had not had chemotherapy or radiation, and were free of known co-morbidities including HIV, chronic viral hepatitis, HNPCC, FAP, and inflammatory bowel disease. Healthy subjects entered the clinic for the study and did not present as a result of co-morbities. Samples were collected from four geographic locations: Toronto (Ontario, Canada), Boston (Massachusetts, USA), Houston (Texas, USA), and Ann Arbor (Michigan, USA). Ninety patients were recruited to the study, thirty of which were designated healthy, thirty with detected adenomas, and thirty with detected carcinomas.

### 16S rRNA Gene Sequence Data Acquisition & Processing

The 16S rRNA gene sequences associated with this study were previously reported (8). Sequence (fastq) and metadata files were downloaded from: http://www.mothur.org/MicrobiomeBiomarkerCRC

The 16S rRNA gene sequences were analyzed as described previously, relying on the mothur software package (v1.37.0) (39, 40). Briefly, the sequences were de-replicated, aligned to the SILVA database (41), screened for chimeras using UCHIME (42), and binned into operational taxonomic units (OTUs) using a 97% similarity threshold. Abundances were normalized for uneven sequencing depth by randomly sub-sampling to 10,000 sequences, as previously reported (23).

### Whole Metagenomic Library Preparation & Sequencing

DNA was extracted from stool samples using the PowerSoil-htp 96 Well Soil DNA Isolation Kit (Mo Bio Laboratories) using an EPMotion 5075 pipetting system. Purified DNA was used to prepare a shotgun sequencing library using the Illumina Nextera XT library preparation kit according to the standard kit protocol, including 12 cycles of limited cycle PCR. The tagmentation time was increased from five minutes to ten minutes to improve DNA fragment length distribution. The library was sequenced using one lane of the Illumina HiSeq4000 platform and yielded 125 bp paired end reads.

### Virus Metagenomic Library Preparation & Sequencing

Genomic DNA was extracted from purified virus-like particles (VLPs) from stool samples, using a modified version of a previously published protocol (29, 31, 43, 44). Briefly, an aliquot of stool (~0.1 g) was resuspended in SM buffer (Crystalgen; Catalog #: 221-179) and vortexed to facilitate resuspension. The resuspended stool was centrifuged to remove major particulate debris then filtered through a 0.22-*μ*m filter to remove smaller contaminants. The filtered supernatant was treated with chloroform for ten minutes with gentle shaking, so as to lyse contaminating cells including bacteria, human, fungi, etc. The exposed genomic DNA from the lysed cells was degraded by treating the samples with 5U of DNase for one hour at 37C. DNase was deactivated by incubating the sample at 75C for ten minutes. The DNA was extracted from the purified virus-like particles (VLPs) using the Wizard PCR Purification Preparation Kit (Promega). Disease classes were staggered across purification runs to prevent run variation as a confounding factor.

As for whole community metagenomes, purified DNA was used to prepare a shotgun sequencing library using the Illumina Nextera XT preparation kit according to the standard kit protocol. The tagmentation time was increased from five minutes to ten minutes to improve DNA fragment length distribution. The PCR cycle number was increased from twelve to eighteen cycles to address the low biomass of the samples, as has been described previously (29). The library was sequenced using one lane of the Illumina HiSeq4000 platform and yielded 125 bp paired end reads.

### Metagenome Quality Control

Both the viral and whole community metagenomic sample sets were subjected to the same quality control procedures. The sequences were obtained as de-multiplexed fastq files and subjected to 5’ and 3’ adapter trimming using the CutAdapt program (v1.9.1) with an error rate of 0.1 and an overlap of 10 (45). The FastX toolkit (v0.0.14) was used to quality trim the reads to a minimum length of 75 bp and a minimum quality score of 30 (46). Reads mapping to the human genome were removed using the DeconSeq algorithm (v0.4.3) and default parameters (47).

### Contig Assembly & Abundance

Contigs were assembled using paired end read files that were purged of sequences without a corresponding pair (e.g. one read removed due to low quality). The Megahit program (v1.0.6) was used to assemble contigs for each sample using a minimum contig length of 1000 bp and iterating assemblies from 21-mers to 101-mers by 20 (48). Contigs from the virus and whole metagenomic sample sets were concatenated within their respective groups. Abundance of the contigs within each sample was calculated by aligning sequences back to the concatenated contig files using the bowtie2 global aligner (v2.2.1), with a 25 bp seed length and an allowance of one mismatch (49). Abundance was corrected for contig reference length and the number of contigs included in each operational genomic unit. Abundance was also corrected for uneven sampling depth by randomly sub-sampling virome and whole metagenomes to 1,000,000 and 500,000 reads, respectively, and by removing samples with fewer total reads than the threshold. Thresholds were set for maximizing sequence information while minimizing numbers of lost samples.

### Operational Genomic Unit Classification

Much like operational taxonomic units (OTUs) are used as an operational definition of similar 16S rRNA gene sequences, we defined closely related bacterial contig sequences as operational genomic units (OGUs) and virus contigs as operational viral units (OVUs) in the absence of taxonomic identity. OGUs and OVUs were defined with the CONCOCT algorithm (v0.4.0) which bins related contigs by similar tetra-mer and co-abundance profiles within samples using a variational Bayesian approach (50). CONCOCT was used with a length threshold of 1000 bp for virus contigs and 2000 bp for bacteria.

### Diversity

Alpha and beta diversity were calculated using the operational viral unit abundance profiles for each sample. Sequences were rarefied to 100,000 sequences. Samples with less than the cutoff were removed from the analysis. Alpha diversity was calculated using the Shannon diversity and richness metrics. Beta diversity was calculated using the Bray-Curtis metric (mean of 25 random sub-sampling iterations), and the statistical significance between the disease state clusters was assessed using an analysis of similarity (ANOSIM) with a post-hoc multivariate Tukey test. All diversity calculations were performed in R using the Vegan package (51).

### Classification Modeling

Classification modeling was performed in R using the Caret package (52). OTU, OVU, and OGU abundance data was preprocessed by removing features (OTUs, OVUs, and OGUs) that were present in less than thirty of the samples. This served both as an effective feature reduction technique and made the calculations computationally feasible. The binary random forest model was trained using the Area Under the receiver operating characteristic Curve (AUC) and the three-class random forest model was trained using the mean AUC. Both were validated using five-fold nested cross validation to prevent over-fitting on the tuning paramters. Each training set was repeated five times, and the model was tuned for mtry values. For consistency and accurate comparison between feature groups (e.g., bacteria, viruses), the sample model parameters were used for each group. The maximum AUC during training was recorded across twenty iterations of each group model to test the significance of the differences between feature set performance. Statistical significance was evaluated using a Wilcoxon test between two categories, or a pairwise Wilcoxon test with Bonferroni corrected p-values when comparing more than two categories.

### Taxonomic Identification of Operational Genomic Units

Operational viral units (OVUs) were taxonomically identified using a reference database consisting of all bacteriophage and eukaryotic virus genomes present in the European Nucleotide Archives. The longest contiguous sequence in each operational genomic unit was used as a representative sequence for classification, as described previously (53). Each representative sequence was aligned to the reference genome database using the tblastx alignment algorithm (v2.2.27) and a strict similarity threshold (e-value < 1e-25) (54). Annotation was interpreted as phage, eukaryotic virus, or unknown. As an additional quality control step, these OVUs were also aligned to the bacterial reference genome set from the European Nucleotide Archives using the blastn algorithm (e-value < 1e-25) and OVUs with similarity to bacterial genomes and not viral genomes were removed from analysis.

### Ecological Network Analysis & Correlations

The ecological network of the bacterial and phage operational genomic units was constructed and analyzed as previously described (34). Briefly, a random forest model was used to predict interactions between bacterial and phage genomic units, and those interactions were recorded in a graph database using *neo4j* graph databasing software (v2.3.1). The degree of phage centrality was quantified using the alpha centrality metric in the igraph CRAN package. A Spearman correlation was performed between model importance and phage centrality scores.

### Phage Replication Style Identification

Phage OVU replication mode was predicted using methods described previously (29, 31, 32). Briefly, we identified temperate OVUs as representative contigs containing at least one of three genomic markers: 1) phage integrase genes, 2) prophage genes from the ACLAME database, or 3) genomic similarity to bacterial reference genomes. Integrase genes were identified in phage OVU representative contigs by aligning the contigs to a reference database of all known phage integrase genes from the Uniprot database (Uniprot search term: “organism:phage gene:int NOT putative”). Prophage genes were identified in the same way, using the ACLAME set of reference prophage genes. In both cases, the blastx algorithm was used with an e-value threshold of 10e-5. Representative contigs were also identified as potential lysogenic phages by having a high genomic similarity to bacterial genomes. To accomplish this, representative phage contigs were aligned to the European Nucleotide Archive bacterial genome reference set using the blastn algorithm (e-value < 10e-25).

## Funding Information

GD Hannigan was supported in part by the Molecular Mechanisms in Microbial Pathogenesis Training Program (T32 AI007528). PD Schloss was supported by funding from the National Institutes of Health (P30DK034933). MT Ruffin was supported by funding from the National Institutes of Health (5U01CA86400). The authors declare no competing interests.

## Acknowledgments

The authors thank the Schloss lab members for their underlying contributions, and the Great Lakes-New England Early Detection Research Network for providing the fecal samples that were used in this study. The authors also thank the participants in the study cohort.

## Supplemental Figure Legends

**Figure S1:**
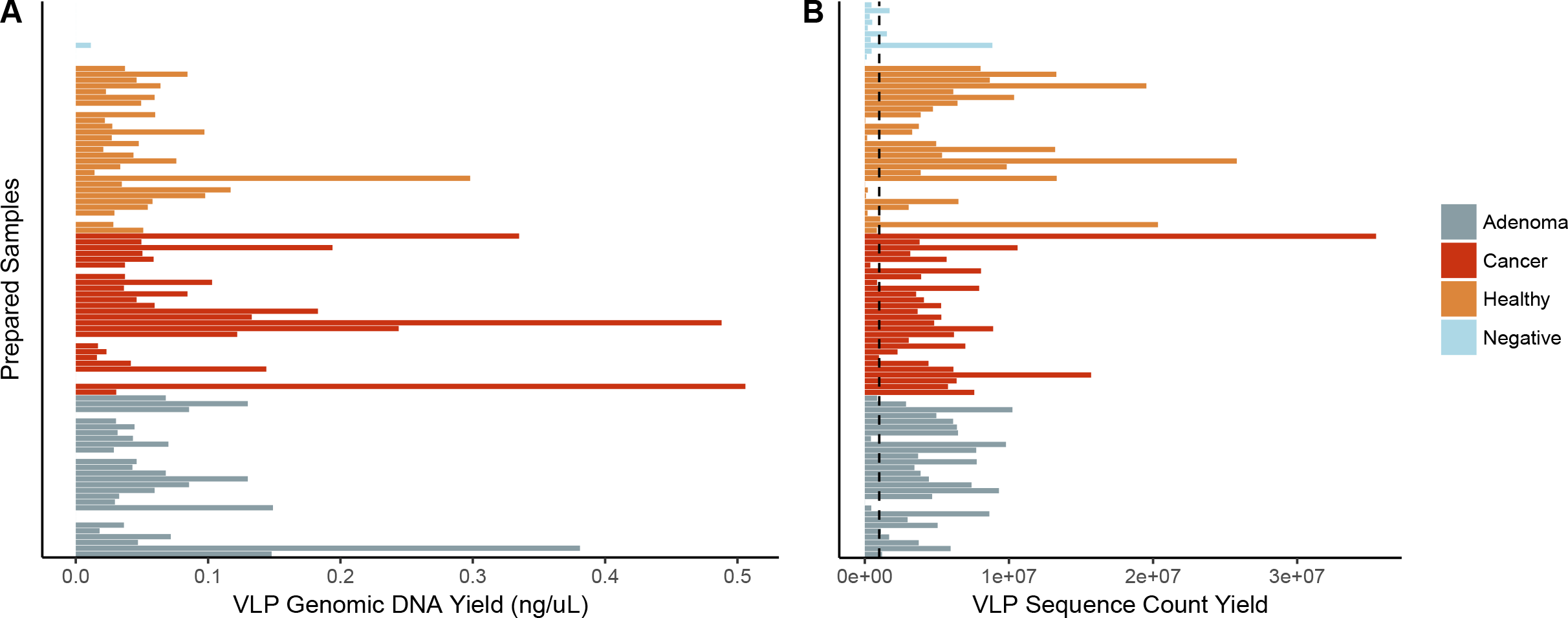
Basic Quality Control Metrics. A) VLP genomic DNA yield from all sequenced samples. Each bar represents a sample which is grouped and colored by its associated disease group or no-DNA negative control. B) Sequence yield following quality control including quality score filtering and human decontamination. Dashed line indicates rarefaction level (10^6^ reads) in which all samples with lower sequence yields less than this level were excluded from downstream analysis. After rarefaction and removal of samples with less than 10^6^ reads, 27 healthy, 28 cancerous, 27 adenomatous, and 3 negative control samples remained.

**Figure S2:**
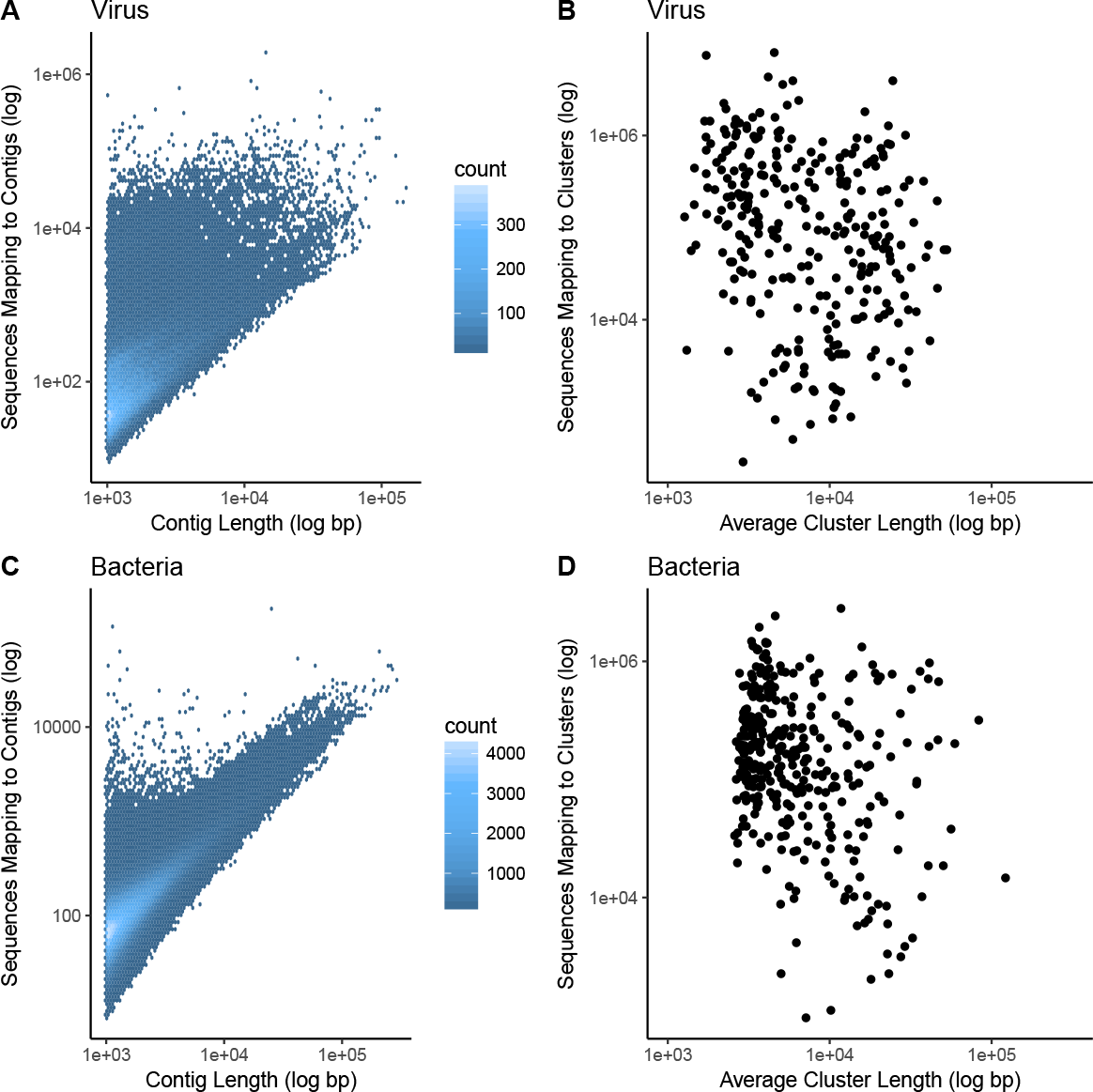
Length and coverage statistics. A) Heated scatter plot demonstrating the distribution of contig coverage (number of sequences mapping to each contig) and contig length for the virus metagenomic sample set. B) Scatter plot illustrating the distribution of operational viral unit (OVU) length and sequence coverage for the virus metagenomic sample set. C) Heated scatter plot demonstrating the distribution of contig coverage and length for the whole metagenomic sample set. D) Scatter plot illustrating the distribution of operational genomic unit (OGU) length and sequence coverage for the whole metagenomic sample set.

**Figure S3:**
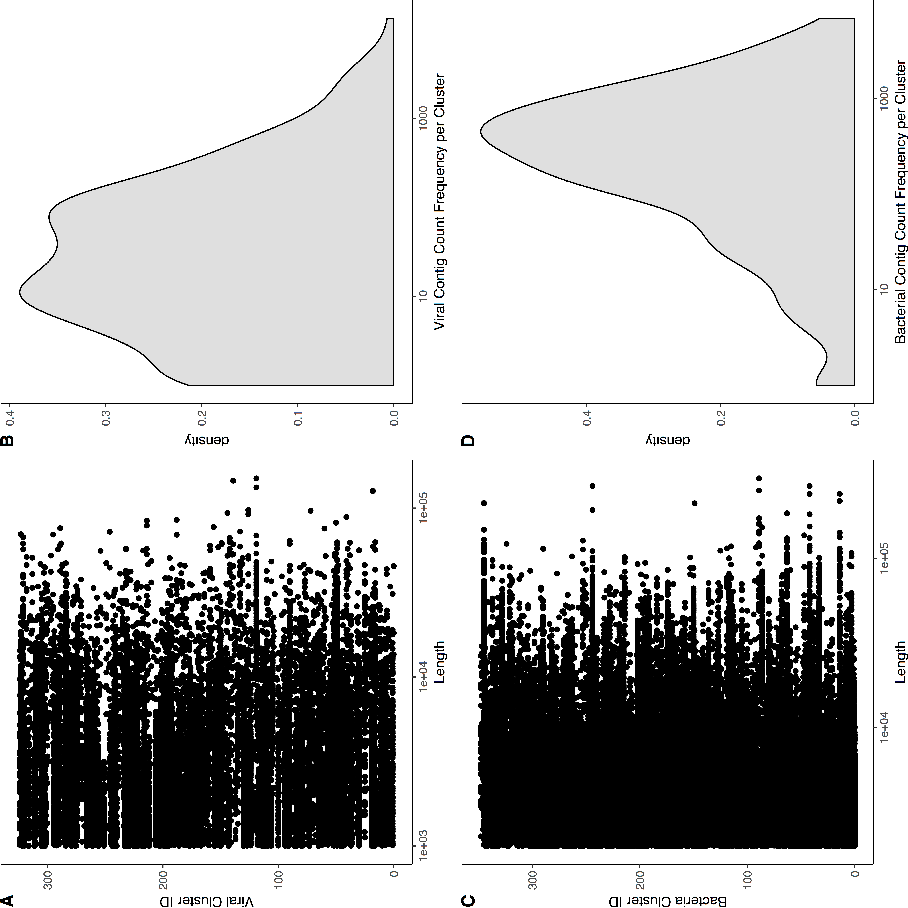
Operational genomic unit composition stats. A) Strip chart demonstrating the length and frequency of contigs within each operational genomic unit of the virome sample set. The y-axis is the operational genomic unit identifier, and x-axis is the length of each contig, and each dot represents a contig found within the specified operational genomic unit. B) Density plot (analogous to histogram) of the number of virome operational genomic units containing the specific number of contigs, as indicated by the x-axis. C-D) Sample plots as panels C and D, but for the whole metagenomic sample set.

**Figure S4:**
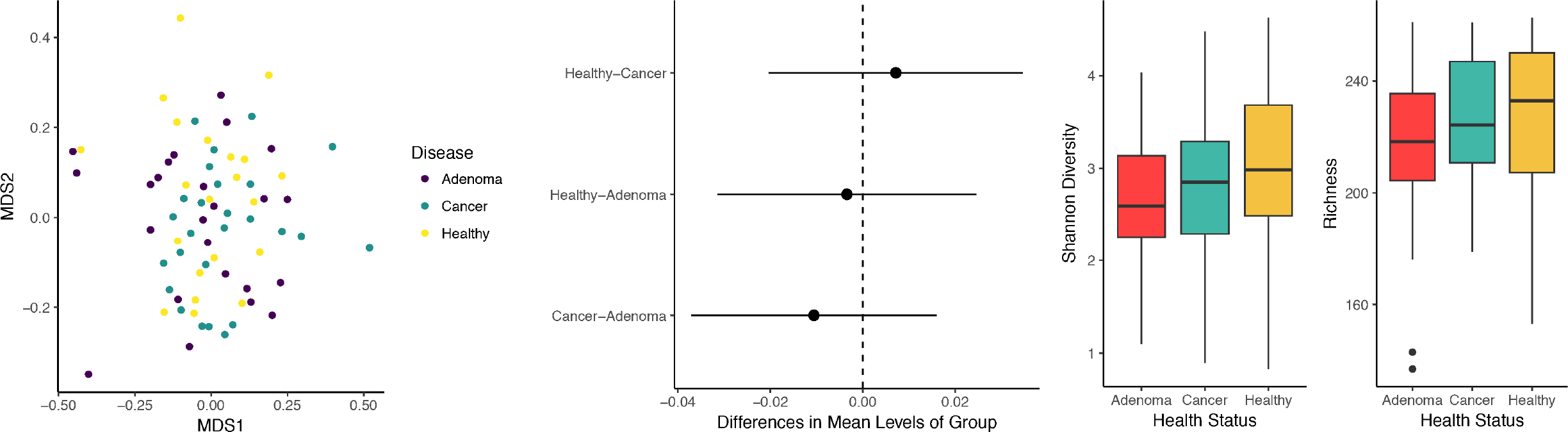
Diversity calculations comparing cancer states of the colorectal virome, based on relative abundance of operational genomic units in each sample. A) NMDS ordination of community samples, colored for cancerous (green), pre-cancerous (red), and healthy (yellow). B) Differences in means between disease group centroids with 95% confidence intervals based on an ANOSIM test with a post hoc multivariate Tukey test. Comparisons (indicated on y-axis) in which the intervals cross the zero mean difference line (dashed line) were not significantly different. C) Shannon diversity and D) richness alpha diversity quantification comparing pre-cancerous (grey), cancerous (red), and healthy (tan) states.

**Figure S5:**
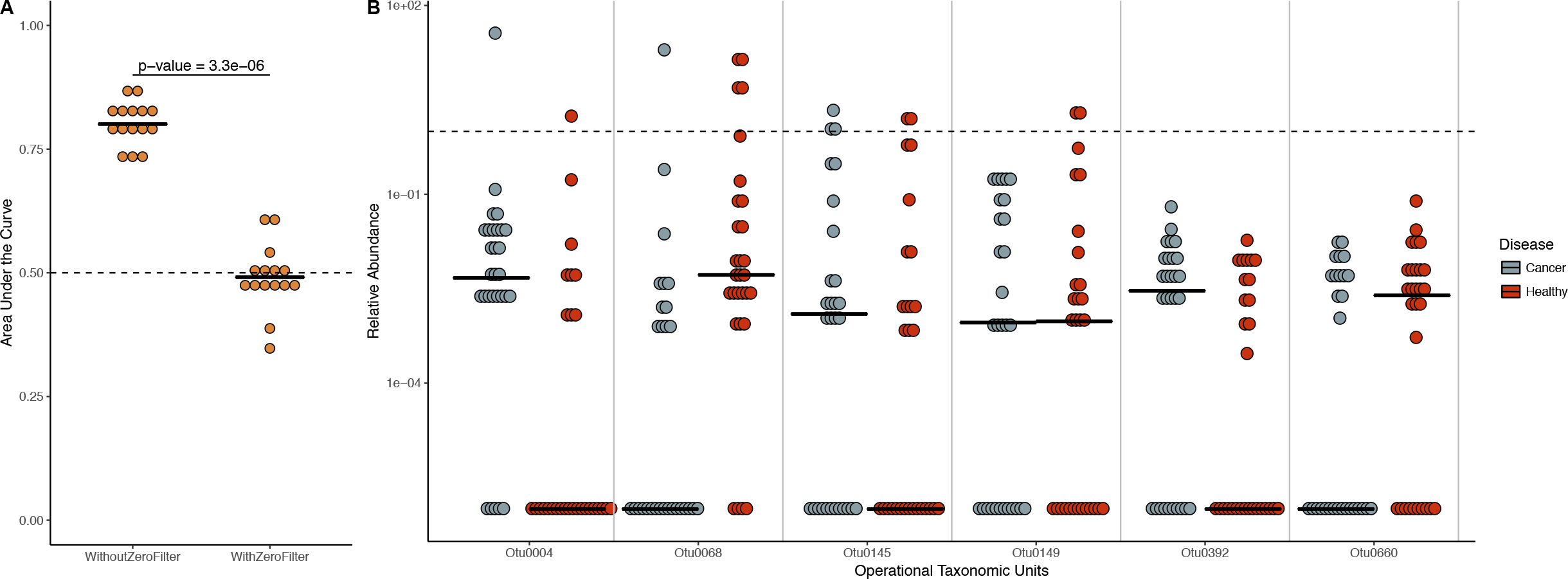
Beta-diversity analysis comparing Bray-Curtis dissimilarity between disease state and negative control community structures that were captured following sequence rarefaction. Differences in means between disease group centroids with 95% confidence intervals based on an ANOSIM test with a post hoc multivariate Tukey test. Comparisons in which the intervals cross the zero mean difference line (dashed line) were not significantly different.

**Figure S6:**
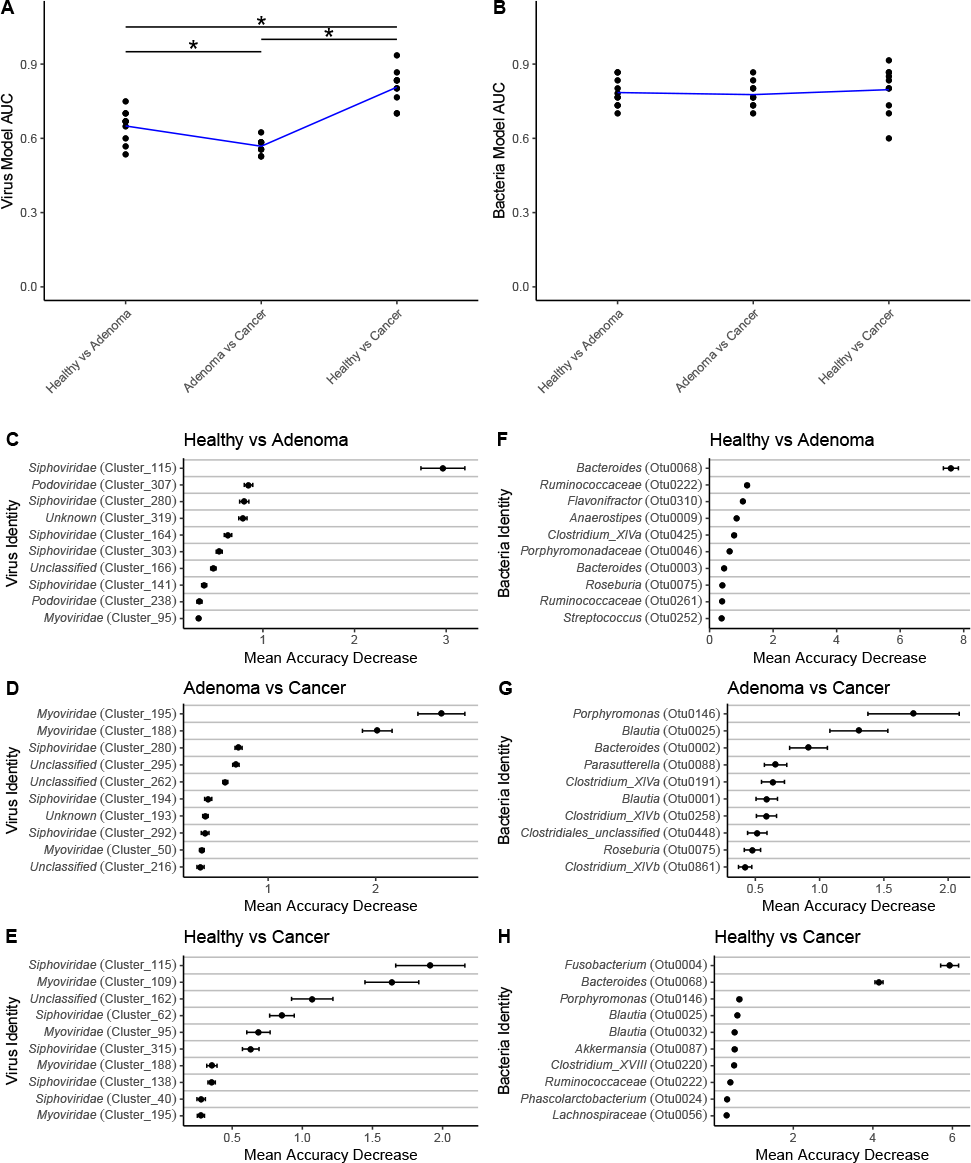
Comparison of bacterial 16S rRNA classification models with and without OTUs whose median relative abundance are greater than zero. A) Classification model performance (measured as area under the curve) for bacteria models using 16S rRNA data both with and without filtering of samples whose median was zero. Significance was calculated using a Wilcoxon rank sum test, and the resulting p-value is shown. The random area under the curve (0.5) is marked with a dashed line. B) Relative abundance of the six bacterial OTUs removed when filtered for OTUs with median relative abundance of zero. OTU relative abundance is separated by healthy (red) and cancerous (grey) samples. Relative abundance of 1% is marked by the dashed line.

**Figure S7:**
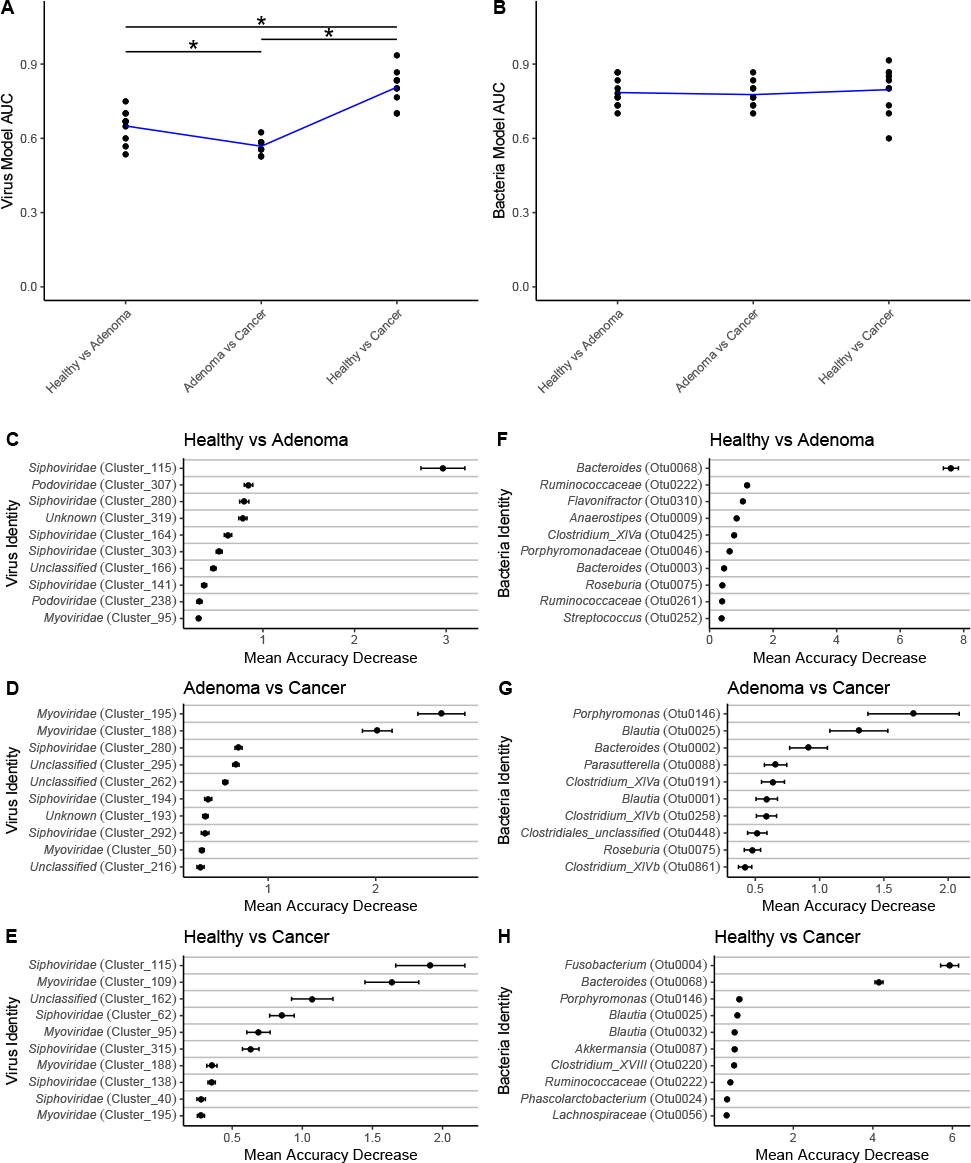
Transition of colorectal cancer importance through disease progression. A) Virus and B) 16S rRNA gene model performance (AUC) when discriminating all binary combinations of disease types. Blue line represents mean performance from multiple random iterations. C-E) Top ten important phage OVUs when classifying each combination of disease state, as measured by the mean decrease in accuracy metric. Mean is represented by a point, and bars represent standard error. Disease comparison is specified in the top left corner of each panel. F-H) Top ten important bacterial 16S rRNA gene OTUs for classifying each disease state combination.

**Figure S8:**
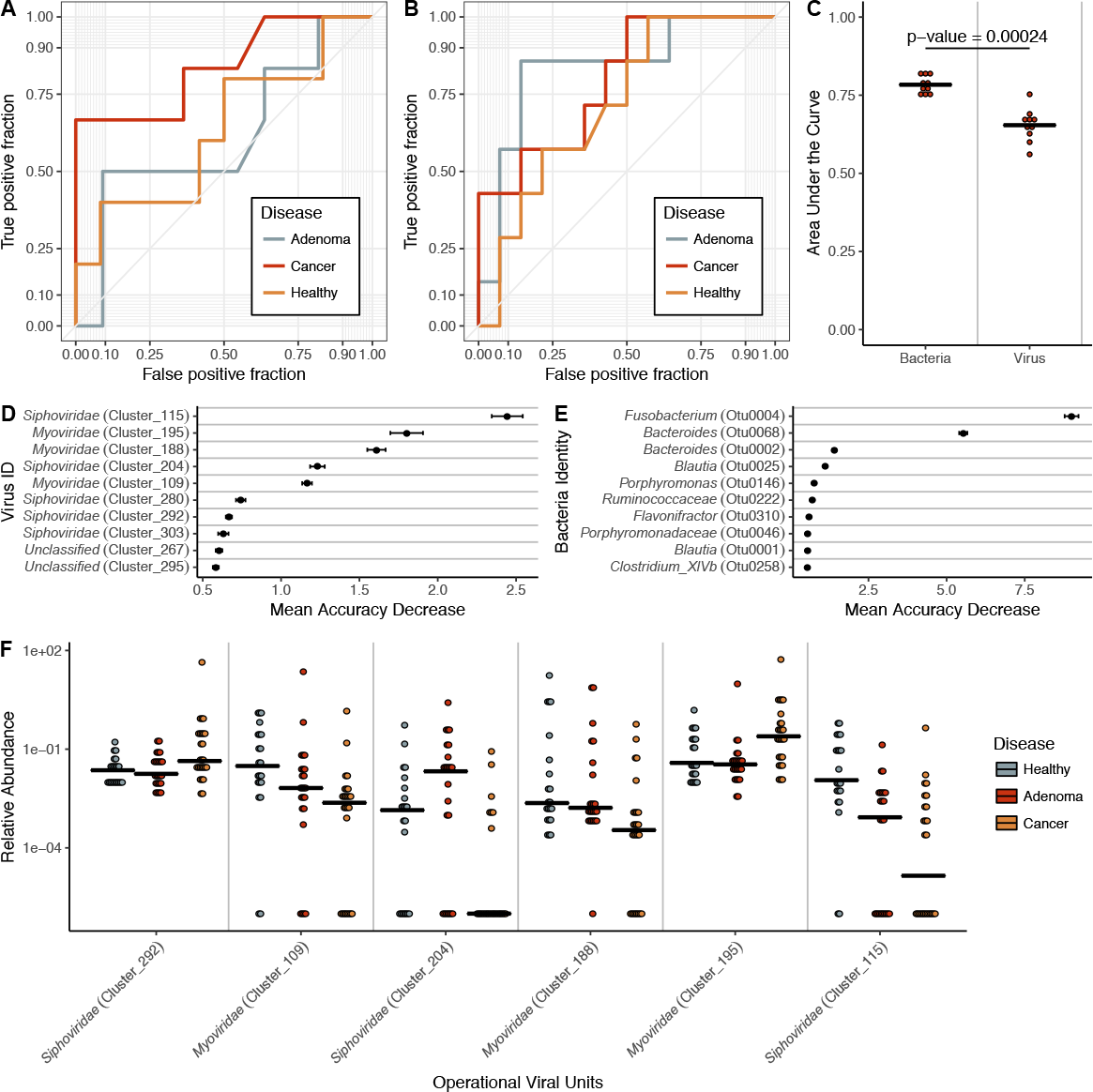
ROC curves from A) virome and B) bacterial 16S three-class random forest models tuned on mean AUC. Each curve represents the ability of the specified class to be classified against the other two classes. C) Quantification of the mean AUC variation for each model based on 10 model iterations. A pairwise Wilcoxon test with a Bonferroni multiple hypothesis correction demonstrated that the models are significantly different (alpha = 0.01). D) Mean decrease in accuracy when virome operational genomic units and E) bacterial 16S OTUs are removed from the respective three-class classification models. Results based on 25 iterations. F) Relative abundance of the six most important virome OVUs in the model, with the most important on the right. Line indicates abundance mean.

